# Biodegradable hyperbranched poly(amine-co-ester)-based polymeric nanoparticles for mRNA delivery

**DOI:** 10.1101/2023.07.20.549815

**Authors:** Gan Liu, Wenqiang Xiang, Miaomiao Guan, Yang Deng

## Abstract

Despite the availability of mRNA vaccines utilizing LNP delivery technology, there remains a pressing need for the development of non-viral mRNA delivery vectors that are both more efficient and safe. we present a novel hyperbranched poly(amine-co-ester) (HBPA) system, catalyzed by immobilized lipase, for efficient *in vitro* and *in vivo* mRNA delivery. By polymerizing four monomers, we successfully synthesized HBPA with a hyperbranched structure, and subsequent modification of the end groups resulted in HBPA-E. Comparative evaluations revealed that HBPA-E outperforms linear PACE and the commercial transfection reagent Lipofectamine MessengerMAX (LipoMM) in terms of intracellular delivery efficiency, while demonstrating lower cytotoxicity. Furthermore, the *in vivo* pulmonary delivery efficiency of HBPA-E was significantly superior to that of LPA-E and the commercial *in vivo* delivery reagent in vivo-JetRNA. Finally, the HBPA-E can be easily dissolved in ethanol, and its mRNA formulation can be employed as a freeze-drying formulation, making it a valuable candidate for future clinical applications of mRNA delivery.

## 1. Introduction

The COVID-19 pandemic has highlighted the critical role of mRNA vaccines in epidemic prevention and control, leading to an unprecedented surge in attention towards mRNA technology[1-3]. Currently, commercial mRNA vaccines utilize non-viral carrier technology based on lipid nanoparticles (LNPs)[4, 5]. These LNPs, formed by combining cationic lipids with mRNA, have demonstrated exceptional efficacy, achieving protection rates exceeding 90% and surpassing other vaccine types. However, LNP technology faces several challenges, including toxic side effects[6-9], limited administration routes[10], difficulties in targeted delivery[11], and stringent storage and transportation requirements, significantly restricting its application range. In recent years, there has been growing interest in cationic polymer-based gene delivery systems[12, 13]. Examples of these systems include widely used polymers such as polyethyleneimine (PEI)[14], polylysine[15, 16], and poly(β-amino ester) (PBAE)[17-19]. Polymeric nanoparticles (PNPs) offer enhanced stability, decreased susceptibility to disintegration, and reduced toxicity compared to other carriers. Moreover, these polymers possess advantages such as cost-effectiveness and straightforward synthesis processes. Saltzman et al. have reported the use of a biodegradable linear cationic poly(amine-co-ester) (PACE) for delivering DNA, mRNA, and siRNA, etc.[20-23]. PACE is synthesized through the polymerization of three small molecular monomers catalyzed by an immobilized lipase. Its high hydrophobicity and low positive charge density contribute to its excellent delivery efficiency and minimal toxicity. Building upon PACE, they have recently developed pulmonary and intranasal mRNA vaccines for COVID-19[24, 25]. However, despite these advancements, PNP-based gene delivery systems have yet to achieve clinical translation primarily due to challenges such as insufficient delivery efficiency, safety concerns, and industrialization issues associated with polymer materials. Hence, the development of novel polymer materials is crucial to construct more promising delivery vectors for clinical applications.

Recently, more and more attention has been paid to gene delivery materials based on hyperbranched polymers, including hyperbranched PEI, polylysine, and PBAE, etc.[26-30]. Hyperbranched polymers offer numerous advantages over their linear counterparts, including enhanced complexation efficiency with genes, improved cellular uptake efficiency, and efficient escape from endosomes[27]. Furthermore, the physical and chemical properties of hyperbranched polymers can undergo significant changes with alterations in their topology and terminal group composition. This versatility suggests that hyperbranched PACE (HBPA) materials may exhibit superior delivery efficiency compared to linear PACE, rendering their development highly valuable.

In this study, we present a novel hyperbranched poly(amine-co-ester) (HBPA) system, catalyzed by immobilized lipase, for efficient *in vitro* and *in vivo* mRNA delivery. By polymerizing four monomers, we successfully synthesized HBPA with a hyperbranched structure, and subsequent modification of the end groups resulted in HBPA-E. Comparative evaluations revealed that HBPA-E outperforms linear PACE (LPA-E) and the commercial transfection reagent Lipofectamine MessengerMAX (LipoMM) in terms of intracellular delivery efficiency, while demonstrating lower cytotoxicity. Furthermore, the *in vivo* pulmonary delivery efficiency of HBPA-E was significantly superior to that of LPA-E and the commercial *in vivo* delivery reagent in vivo-JetRNA. An intriguing aspect of HBPA-E synthesis, facilitated by immobilized enzyme catalysis, is the absence of any cross-linking between polymers. This unique feature differentiates it from conventional branched polymerization methods that often lead to undesired cross-linking. Notably, HBPA-E exhibits solubility in ethanol, unlike LPA-E. This characteristic enables the formulation of polymer nanoparticles (PNPs) without the need for other potentially toxic solvents, thereby enhancing its potential for clinical translation.

## 2. Results and discussion

### 2.1 Synthesis and Characterization of hyperbranched polymers

Similar to the synthesis method of PACE reported by Saltzman et al.[20], we utilized an immobilized lipase to catalyze the copolymerization of four monomers to synthesize hyperbranched polymers. In this process, we introduced triethanolamine, which contains three hydroxyl groups, to promote the formation of branched structures. Unlike conventional branched polymerization methods that often result in cross-linking, our polymers did not undergo cross-linking at the end of the reaction. By adjusting the amounts of N-methyldiethanolamine (M) and triethanolamine (T) among the four monomers, we obtained linear polymer LPA and hyperbranched polymer HBPA with varying degrees of branching. The ^1^H-NMR spectra of these polymers, as shown in Fig. 2A, revealed characteristic peaks corresponding to different monomer units. Peaks a, b, and f represented unit PDL (P), while peaks a, b, and c corresponded to unit SA (S). Peaks e and g indicated the presence of unit MDEA (M), and peaks h and i represented unit TEA (T). The intensities of the peaks associated with P and S units were found to be similar among different HBPA, indicating the almost same composition of these units. Whereas, the signals from M (e and g) gradually decreased with decreasing M content, and the signals from T (h and i) increased with increasing T content. These results demonstrate that the chemical composition of the polymers can be precisely controlled by adjusting the types and proportions of the monomers.

**Fig. 1.**
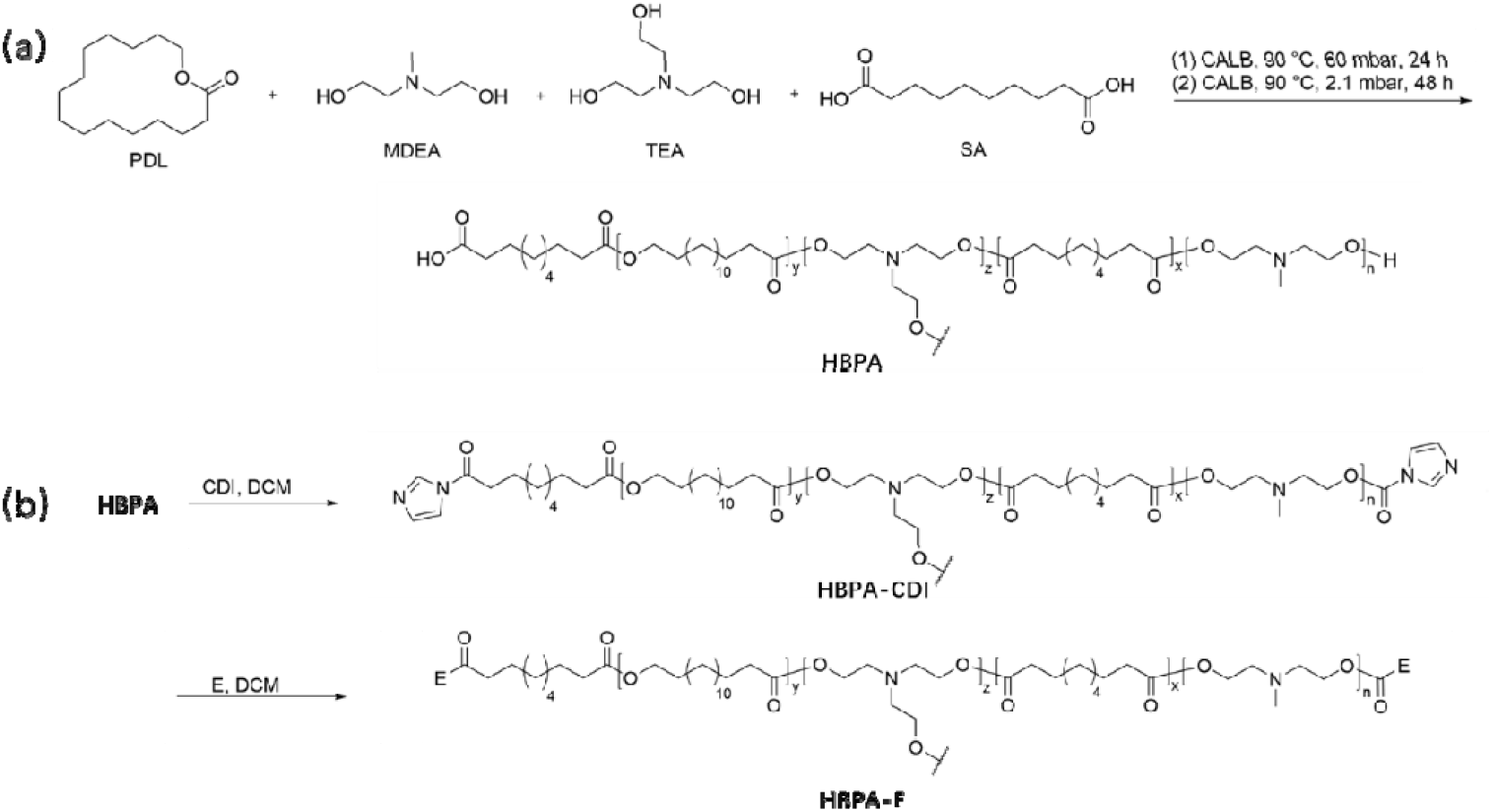
Synthesis of hyperbranched poly(amine-co-ester) (HBPA) tetrapolymers. (a) Two-stage process for copolymerization of lactone with MDEA, TEA and SA catalyzed by an immobilized lipase. (b) Synthesis of HBPA-E through end group modification of HBPA with E monomers.

**Fig. 2.**
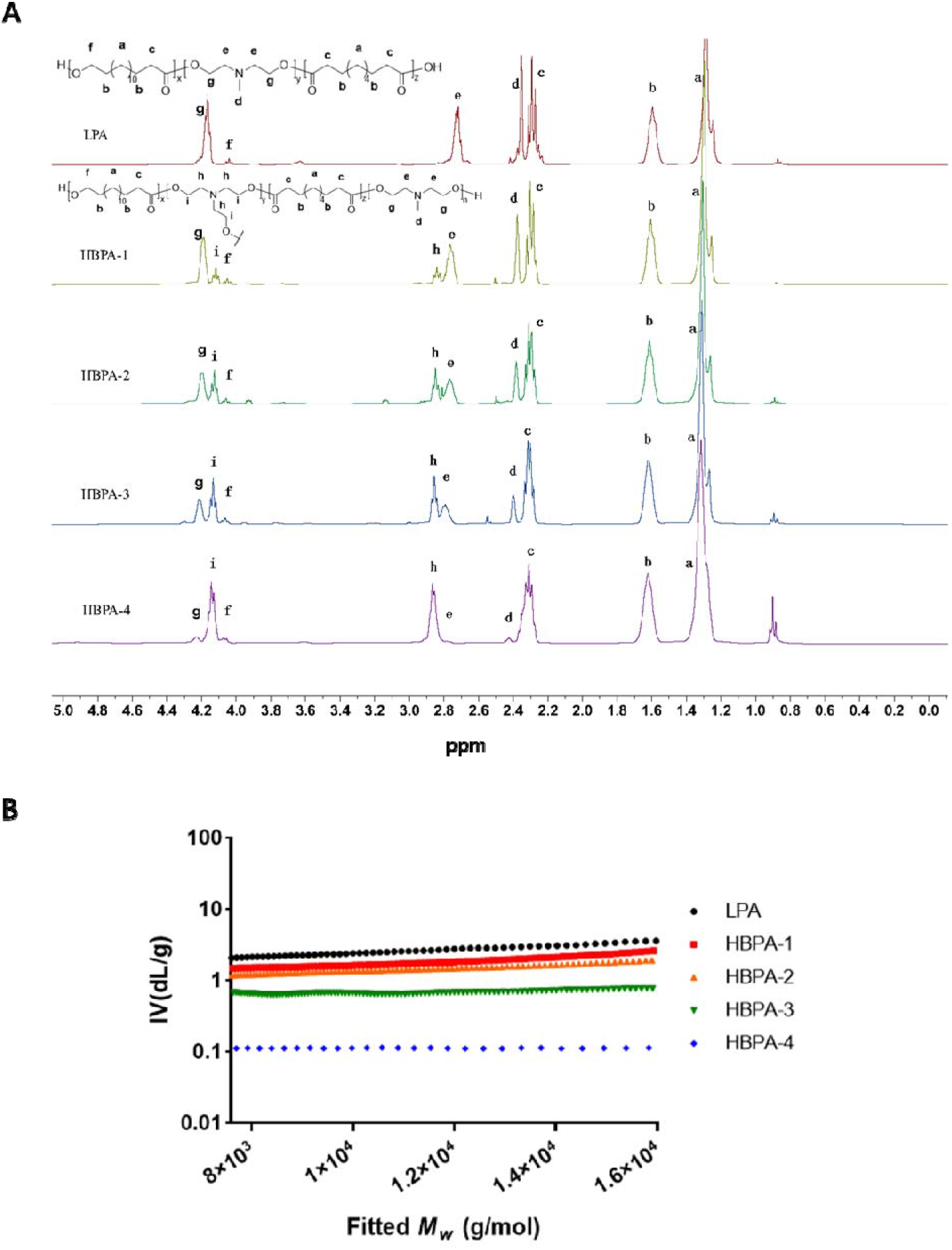
Characterization of the highly branched HBPA with different branched structures. (A) ^1^H NMR spectra of the 5 versions of HBPAs and LPA. As the feed ratio of TEA/MDEA increased, the TEA units in HBPA increased, whereas the MDEA units decreased correspondingly. (B) MH plots of the HBPA and the corresponding LPA. Increases in the feed ratio of TEA/MDEA resulted in a sequential decrease of a values of the LPA and HBPAs.

Furthermore, we conducted structural characterization of the different polymers using gel permeation chromatography (GPC). The degree of branching, indicated by the α value, was calculated based on the MH plot as shown in Fig. 2B. We observed a gradual decrease in the α value from 0.51 for HBPA-1 to 0.31 for HBPA-4, indicating an increasing degree of branching. This finding aligns with the results obtained from the ^1^H-NMR analysis. Thus, it can be concluded that the incorporation of triethanolamine (T) into the polymer synthesis process promotes branching in the polymer structure, and the degree of branching increases with higher T content.

Table 1 presents the properties of E14-modified HBPA (HBPA-E14). It is evident that the component proportions align with the proportions of the added monomers. The average molecular weight (Mw) is approximately 20 k, and the polydispersity index (PDI) is around 3. Notably, HBPA-E14 exhibits excellent solubility in ethanol (>25 mg/mL), which is in stark contrast to LPA-E14. This observation suggests that the branched structure of the polymer significantly impacts its properties.

**Table 1.** Characterization of selected HBPA-E14 and LPA-E14.

| Name | P/S/M/T<br>(feed molar<br>ratio) | P/S/M/T (unit<br>molar ratio)<sup>a</sup> | $M_n^b$ | $M_w^b$ | PDI<sup>b</sup> | Solubility<br>in alcohol<br>( mg/mL<br>) |
| --- | --- | --- | --- | --- | --- | --- |
| LPA-E14 | 1: 9: 9: 0 | 1: 10: 9.11: 0 | 8769 | 17275 | 1.97 | <10 |
| HBPA-E14-1 | 1: 9: 6.2: 1 | 1: 9.9: 4.9: 1.0 | 5457 | 18172 | 3.33 | >25 |
| HBPA-E14-2 | 1: 9: 6: 2 | 1: 9.0: 4.5: 1.9 | 6548 | 21084 | 3.22 | >25 |
| HBPA-E14-3 | 1: 9: 4.5: 3 | 1: 8.9: 4.1: 2.9 | 7534 | 21472 | 2.85 | >25 |
| HBPA-E14-4 | 1: 9: 3: 4 | 1: 8.5: 2.9: 3.6 | 7473 | 22045 | 2.95 | >25 |

<sup>a</sup>Measured by <sup>1</sup>H NMR spectroscopy. <sup>b</sup>Measured by gel permeation chromatography (GPC) with DMF as solvent.

### 2.2 mRNA transfection with hyperbranched polymers

We subsequently evaluated the cellular transfection efficiency of the hyperbranched polymer HBPA-E14 in delivering Luciferase mRNA, and compared it with the commercial transfection reagent LipoMM and the linear polymer LPA-E14. As depicted in Figure 3A, at a polymer-to-mRNA weight ratio of 50:1, HBPA-E14-1 demonstrated significantly higher mRNA transfection efficiency in A549 cells compared to LipoMM, with nearly a three-fold increase, surpassing the performance of linear LPA-E14. This finding highlights the superior efficacy of the hyperbranched polymer over the linear counterpart, as well as its superiority over the most efficient commercial transfection reagent.

**Fig. 3.**
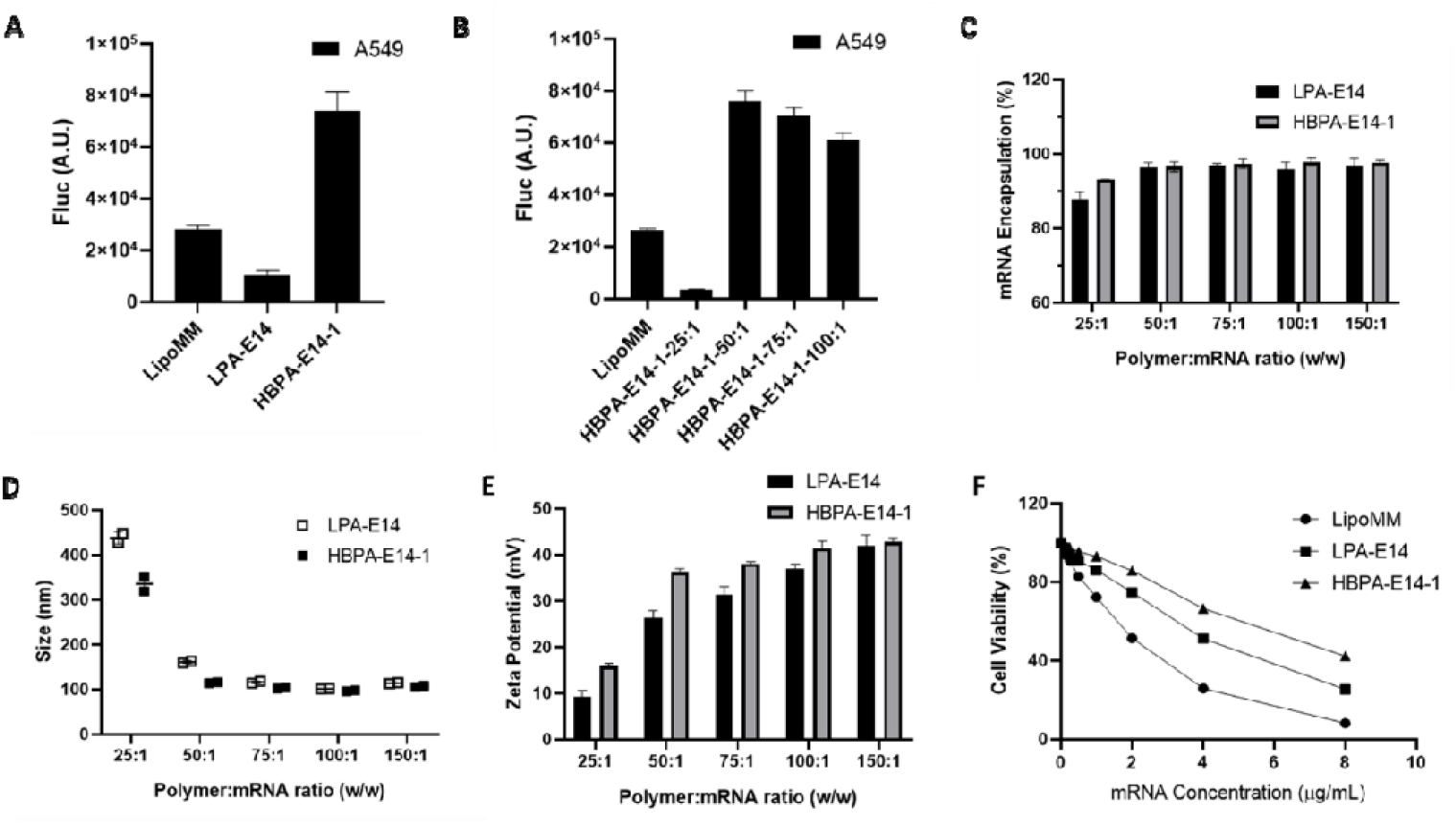
HBPA-E14 show higher mRNA transfection efficiency compared to LPA-E14. (A) Transfection effciency of HBPA-E14-1 compared with LPA-E14 on A549 cells. A.U., Arbitrary Unit. (B) The effect of weight ratio of HBPA-E14-1 to mRNA on transfection efficiency on A549 cells. (C) mRNA encapsulation efficiency of HBPA-E14-1 compared to LPA-E14. (D) Size and (E) Zeta potential of HBPA-E14-1 compared to LPA-E14. (F) Cytotoxicity of LipoMM, LPA-E14 and HBPA-E14-1 on A549 cells.

Furthermore, we investigated the transfection efficiency of HBPA-E14-1/mRNA complexes at different weight ratios of polymer to mRNA (25:1, 50:1, 75:1, and 100:1) in A549 cells. As illustrated in Figure 3B, complexes with weight ratios ranging from 50:1 to 100:1 exhibited comparable and significantly higher transfection efficiency compared to LipoMM. However, the complexes formed with a weight ratio of 25:1 showed negligible transfection. These results suggest that when the weight ratio reaches 50:1 or higher, the polymer and mRNA are adequately complexed, resulting in efficient transfection. Conversely, at a weight ratio of 25:1, the complexation is insufficient to achieve detectable transfection.

We conducted further evaluations of the mRNA encapsulation efficiency, size, and Zeta potential of HBPA-E14-1/mRNA complexes at various weight ratios, comparing them with LPA-E14/mRNA complex. Fig. 3C presents the mRNA encapsulation efficiencies of HBPA-E14-1 and LPA-E14 at different weight ratios. The results indicated that both polymers achieved encapsulation efficiencies exceeding 95% when the weight ratio ranged from 50:1 to 150:1. However, at a weight ratio of 25:1, the mRNA encapsulation efficiency of HBPA-E14-1 and LPA-E14 was only 93% and 87.8%, respectively. These values were significantly lower than the higher weight ratios, indicating inadequate complexation between the polymer and mRNA at this weight ratio, aligning with the previous findings on cellular transfection. Additionally, at the weight ratio of 25:1, HBPA-E14-1 exhibited higher mRNA encapsulation efficiency compared to LPA-E14, suggesting stronger complexation of HBPA-E14-1 with mRNA than LPA-E14.

The size and Zeta potential of the complexes, as depicted in Fig. 3D and 3E, displayed similar trends. When the weight ratio was 50:1 to 150:1, the complexes formed by both polymers had a size of approximately 100 nm and a Zeta potential of around 30 mV. However, at the weight ratio of 25:1, the particle size of the two complexes was notably higher than at other weight ratios, while the Zeta potential was significantly lower. Furthermore, at the weight ratio of 25:1, HBPA-E14-1 exhibited smaller particle size and higher Zeta potential compared to LPA-E14. These results collectively indicate that HBPA-E14-1 demonstrates stronger complexation efficiency with mRNA compared to LPA-E14. Moreover, both polymers can form stable complexes with mRNA when the weight ratio reaches 50:1 or higher.

Additionally, we carried out cytotoxicity tests on various polymer-mRNA complexes. As shown in Fig. 3F, the control LipoMM complex exhibited significant cytotoxicity, resulting in only a 50% cell survival rate at an mRNA concentration of 2 μg/mL. Moreover, the toxicity increased noticeably with higher mRNA concentrations. In contrast, the linear LPA-E14 complex displayed reduced toxicity compared to the control, although some cytotoxicity was still observed, resulting in a 70% cell survival rate at an mRNA concentration of 2 μg/mL. Remarkably, the hyperbranched HBPA-E14-1 complex demonstrated even lower toxicity, with a cell survival rate of over 80% at the mRNA concentration of 2 μg/mL. These findings indicate that the hyperbranched HBPA-E14-1 complex exhibits higher biocompatibility than both the commercial transfection reagent and the linear LPA-E14 complex, making it more promising for future *in vivo* applications.

We further conducted experiments to evaluate the mRNA transfection efficiency of HBPA-E14 polymers with varying branching degrees in different cell lines, comparing them with LipoMM and LPA-E14. The results, depicted in Fig. 4A-4D, clearly demonstrate that across A549, HEK293T, HeLa, and HT22 cells, all four variants of HBPA-E14 exhibited significantly higher mRNA transfection efficiency compared to the commercial control, LipoMM. Specifically, the mRNA transfection efficiency of HBPA-E14 polymers was 1-5 times higher than that of LipoMM and substantially surpassed the transfection efficiency of linear LPA-E14. Notably, HBPA-E14-2 displayed the highest transfection efficiency in A549, HeLa, and HT22 cells, while HBPA-E14-1 exhibited the highest efficiency in HEK293T cells. These findings strongly suggest that hyperbranched polymers with different branching degrees are more effective for mRNA transfection in various cell lines compared to linear polymers and also outperform the most efficient commercial transfection reagents. Moreover, they demonstrate their universal applicability across different cell types.

**Fig. 4.**
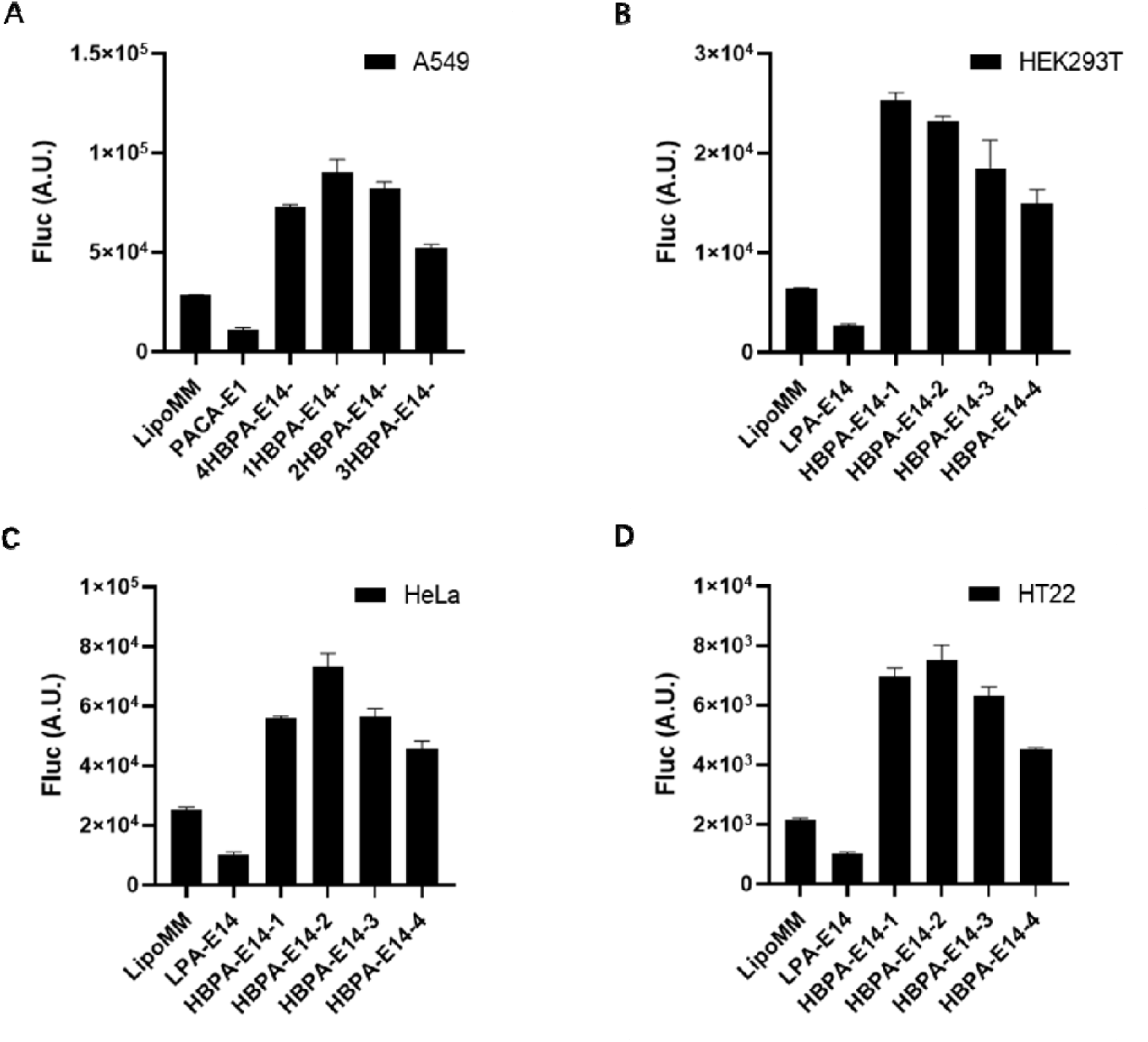
HBPA-E show higher mRNA transfection efficiency compared to LPA-E14 over diverse cell types. (A) A549, (B) HEK293T, (C) HeLa and (D) HT22 cells.

### 2.3 Effectiveness of mRNA delivery *in vivo*

In our final experiment, we assessed the *in vivo* mRNA delivery efficacy of HBPA-E14-1 (HBPA) through pulmonary administration. We compared the efficiency of different groups, including LPA/LPA-PEG, HBPA, HBPA/HBPA-PEG, and HBPA/HBPA-PEG freeze-dried formulations, for delivering luciferase mRNA to the lungs. As controls, we included a blank Control group and a commercial in vivo-jetRNA group. Fig. 5 illustrated no fluorescence signal was detected in the Control group, while the luciferase expression efficiency of the in vivo-jetRNA group was defined as 1. The luciferase expression efficiency of LPA/LPA-PEG was comparable to that of in vivo-jetRNA. Notably, the luciferase expression efficiency of HBPA, HBPA/HBPA-PEG, and HBPA/HBPA-PEG freeze-dried agents was 3.5, 22, and 45 times as that of in vivo-jetRNA, respectively. These results clearly demonstrate that the hyperbranched polymer HBPA exhibits superior delivery efficiency compared to the linear polymer LPA. Furthermore, when combined with PEG, the delivery efficiency of HBPA *in vivo* is further enhanced. It is worth noting that the properties of the hyperbranched polymer complex after lyophilization can be maintained or even improved, facilitating transportation, storage, and practical utilization of the formulation. Therefore, this hyperbranched polymer holds immense potential for application in the field of mRNA delivery.

**Fig. 5.**
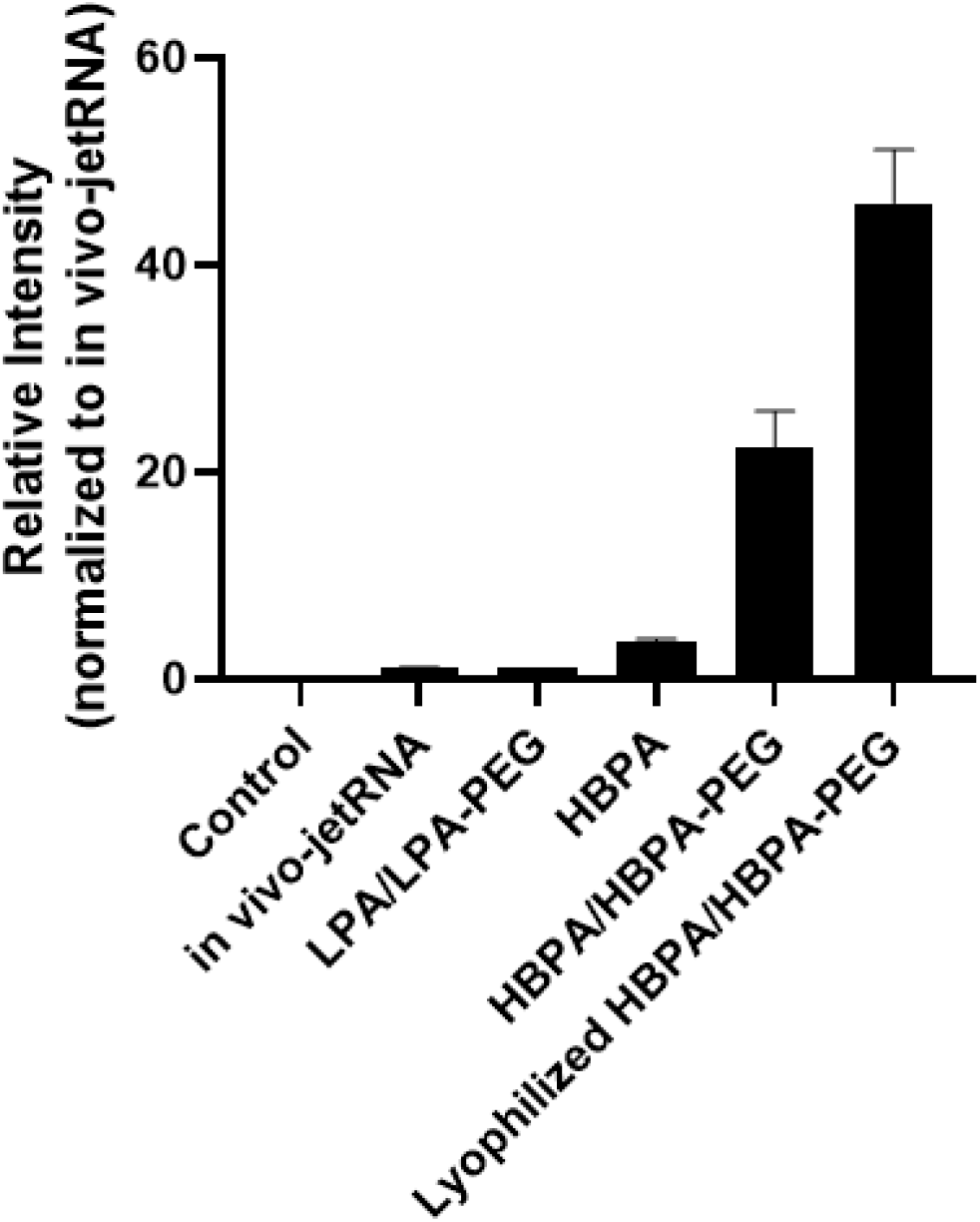
Relative luciferase protein expression in lung tissue after delivery of FLuc mRNA with LPA/LPA-PEG, HBPA, HBPA/HBPA-PEG and Lyophilized HBPA/HBPA-PEG.

## 3 Conclusion

We have successfully developed a hyperbranched cationic polyester, HBPA-E, through immobilized enzyme catalysis, for efficient mRNA delivery both *in vitro* and *in vivo*. This hyperbranched material demonstrates superior cell delivery efficiency and lower cytotoxicity compared to linear LPA-E and the commercial transfection reagent, LipoMM. Furthermore, the *in vivo* delivery efficiency of HBPA-E surpasses that of linear LPA-E and the commercial in vivo-jetRNA. The advantages of HBPA-E extend beyond its delivery efficiency. The material can be easily dissolved in ethanol, and its production process can be well controlled. Moreover, the complex formed between HBPA-E and mRNA can be employed as a freeze-drying formulation, enhancing its stability and usability. Considering the similarities between different gene materials, it is plausible to anticipate the use of HBPA-E for the delivery of other genes, thereby highlighting its substantial potential for clinical translation and application. These findings underscore the promising clinical prospects of HBPA-E as an effective and safe mRNA delivery system. Its enhanced delivery efficiency, low cytotoxicity, ease of production, and potential applicability to various gene materials position HBPA-E as a valuable candidate for future clinical transformations and gene delivery applications.

## Acknowledgements

Special thanks to Professor Dezhong Zhou at Xi’an Jiaotong University, School of Chemical Engineering and Technology for assistance with GPC data analysis.

## Author contributions

YD and LG conceived the idea. GL, WQX, MMG and YD designed the whole experimental project and carried out the synthesis, characterization and data analysis. GL, WQX, MMG and YD wrote the manuscript and provided revisions.

## Supplementary Materials

Materials and Methods

## Materials and Methods

### 1. materials

The following chemicals and reagents were acquired from Aladdin Scientific (Shanghai, CN): pentadecanolactone (PDL), sebacic acid (SA), N-methyldiethanolamine (MDEA), triethanolamine (TEA), phenyl ether, hexane, 1,1’-carbonyldiimidazole (CDI), 1,3-Diamino-2-propanol, dichloromethane, chloroform-d, 1,2-Diaminopropane, dithiothreitol (DTT) and 1,3-Diamino-2-propanol. The immobilized Candida antarctica lipase B (CALB) supported on acrylic resin (Novozym 435) was obtained from Sigma Aldrich. mPEG_2k_-NH_2_ was purchased from Bide Pharmatech (Shanghai, CN). Lipofectamine MessengerMAX was procured from ThermoFisher Scientific. Firefly Luciferase mRNA (N1-Me-Pseudo UTP) was purchased from Vazyme Biotech Co., Ltd. Fetal bovine serum (FBS) was sourced from Sigma-Aldrich (Merck), while DMEM was obtained from Corning, and Trypsin (0.25% Trypsin-EDTA 1X) was obtained from ThermoFisher Scientific. A549, HEK293T, HeLa, and HT22 cell lines were obtained from ATCC. D-Luciferin was purchased from Macklin. C57BL/6 mice were acquired from Guangzhou Ruige Biotechnology Co., LTD.

### 2. Synthesis of polymers

The copolymerization of PDL (P) with SA (S), MDEA (M) and TEA (T) was performed in diphenyl ether solution using a parallel synthesizer connected to a vacuum line with the vacuum (±0.2 mmHg) controlled by a digital vacuum regulator. In a typical experiment, reaction mixtures were prepared, which contained four monomers (P, S, M and T, monomers molar ratio as Table 1), Novozym 435 catalyst (10 wt% versus total monomer), and diphenyl ether solvent (200 wt% versus total monomer). The copolymerization reactions were carried out at a constant temperature in two stages: first-stage oligomerization, followed by second-stage polymerization. During the first-stage reaction, the reaction mixtures were stirred at 90 □ under 60 mBar of argon gas for 24 hours, after which the reaction pressure was reduced to 2.1 mBar and the reactions were continued for a further 24 h. The polymer products were isolated and purified according to the following procedures.

The crude product mixtures were first diluted with a small amount of dichloromethane, and then filtered to remove catalyst particles. The filtrates were mixed with hexane to cause the precipitation of the polymers. The precipitated polymers were then washed several times with fresh hexane to extract and remove the residual diphenyl ether solvent from the polymers. Evaporation and complete removal of the solvent from the precipitates under high vacuum yielded the purified polymers HBPA.

To modify HBPA with 1,3-Diamino-2-propanol (E14), excessive amount of CDI were added into the solution of HBPA polymers in dry dichloromethane, which was stirred for 6h at room temperature. Then excessive amount of E14 was added into the above solution, and then continue to react overnight. After completion of reaction, the mixture was washed three times with twice volume of deionized water until aqueous phase was neutral, followed by evaporation of DCM under vacuum to obtain HBPA-E14.

To synthesize PEGylated HBPA (HBPA-PEG), CDI at a molar ratio of 1:1 was added into a solution of HBPA in dry dichloromethane, the mixture was stirred for 6 h at room temperature, followed by the same ratio of mPEG-NH_2_ and stirred for further 1 day. Extra 10 equivalent of CDI was added in the above solution. After another 6 h, the mixture was treated with excessive ethanolamine and stirred overnight. Then the mixture was washed three times with twice volume of deionized water until aqueous phase was neutral, followed by evaporation of DCM under vacuum to obtain PEGylated polymers.

### 3. Characterization of polymers

1H-NMR spectra were recorded using TMS as the internal standard in CDCl_3_ with a Bruker BioSpin GmbH spectrometer at 400 MHz, respectively. Molecular weights (M_w_ and M_n_) and PDI values of polymers were measured by gel permeation chromatography (GPC). Base polymers in Fig. 2 were analyzed using an Agilent 1260 Infinity GPC system equipped with a refractive index detector, a viscometer detector, and a dual-angle light-scattering detector (LS 15° and LS 90°). GPC columns (PolarGel-M, 7.5 mm × 300 mm; two in series) were eluted with DMF and 0.1% LiBr at a flow rate of 1 ml/min at 60 °C. GPC columns were calibrated with linear poly(methyl methacrylate) standards. To analyze the molecular weights of the base polymers, 20 μl of the reaction solution was taken at different time points, diluted with 1 ml of DMF, filtered through a 0.2-μm filter, and then measured by GPC. Base polymers in Table 1 were analyzed using the waters 1515 chromatography and the 2414 refractive index (RI) detector at 35 □. The flow phase is a DMF with 0.1% LiBr, velocity 1 ml/min, standard of linear polymethyl methacrylate.

### 4. Preparation and characterization of polyplexes

For *in vitro* experiment, various amounts of polymer solution (40 mg/mL in ethanol for HBPA-E and in DMSO for LPA) was first diluted in 50 μL sodium acetate buffer. After brief vortexing, the polymer solution was mixed with 1 μg mRNA diluted in 50 μL sodium acetate buffer and vortexed again to prepare polyplexes with polymer:mRNA weight ratio as 25:1, 50:1, 75:1, 100:1 and 150:1. The polymer:mRNA mixture was incubated at room temperature for 10 min before use. For *in vivo* experiments, a solution at 100 μg mRNA/mL in sodium acetate buffer was prepared by the same method. Unless specifified, polymer:mRNA polyplexes were prepared at a 50:1 polymer:mRNA weight ratio in 9 mM sodium acetate buffer (pH 4.9) for *in vitro and in vivo* experiments.

HBPA-E14-1/HBPA-PEG-1/mRNA polyplex was prepared by similar method. HBPA-E14-1 and HBPA-PEG-1 were dissolved together in ethanol and then diluted into sodium acetate buffer. Then the blended polymer solution was mixed with mRNA solution to get polyplexes with HBPA-E14-1:HBPA-PEG-1:mRNA weight ratio as 50:5.6:1. To prepare lyophilized formulation of polyplex, trehalose aqueous solutions (500 mg/mL) were added to the polyplex suspension at a certain volume ratio to obtain final trehalose concentrations of 15 mg/mL. The mixtures were then snap frozen in ultra-low temperature refrigerator and lyophilized for 1 days. The lyophilized polyplexes was stored at 4 °C and resuspended in RNase-free water for use when needed.

The hydrodynamic diameter of the polyplexes was measured by Dynamic Light Scattering (DLS) using a Malvern Zetasizer Lab (Malvern Instruments, UK), after dilution of polyplexes in ultrapure water at a concentration of 2 μg/mL of mRNA. To measure zeta potential, the same solution was loaded into a disposable capillary cell and analyzed on a Malvern Zetasizer Lab.

Encapsulation efficiency (EE) of mRNA in the polyplexes was measured using the Quant-IT RiboGreen RNA kit (Invitrogen) according to manufacturer instructions. By using RiboGreen assay measures the amount of free mRNA in solution and the amount of total mRNA after samples were treated with 0.5% Triton X-100, to calculate encapsulation efficiency.

### 5. *In vitro* mRNA transfection efficiency study

After obtaining polyplexes of different HBPA-E with mRNA, the transfection efficiency was tested in different cell lines. A549, HEK293T, HeLa and HT22 cells were seeded in 48-well cell culture plates (2.5×10^4^/ well). The cells were cultured in DMEM containing 5% CO_2_ at 37 □ for 12 h, followed by DMEM medium (250 μL/ well). Samples to be tested and positive controls were added and cultured for 24 h, respectively, and samples were added according to the final 2 μg/mL mRNA/well. For positive control of mRNA transfection, Lipofectamine MessengerMAX (LipoMM) was used. After transfection, the cells were aspirated out of the medium in the well plates and then placed in a refrigerator at -80 ° C for 10 minutes. The well plates were carefully extracted and stored at 4 °C. Subsequently, 40 μL of lysate was transferred to each well of a 48-well plate, ensuring complete coverage of the bottom surface, and left to incubate for 5 minutes. Simultaneously, 280 μL of the test solution was added to another 48-well plate. Then, 160 μL of the resulting mixture was carefully transferred to a black microplate reader plate. Finally, a precise amount of 40 μL of D-Luciferin was added to each well using a pipette. After 2 minutes at room temperature, the reaction was put into a microplate reader to test the luminescence at 560 nm.

### 6. *In vitro* cytotoxicity study

The cytotoxicity of the HBPA-E/mRNA polyplex was tested in A549 cells. A549 cells were seeded in 96-well cell culture plate (1×10^4^/well). After cell culture for 12 h, the cells were attached to the wall, replaced by DMEM medium, and the mRNA polyplexes of LPA-E14, HBPA-E14-1 and positive control LipoMM were added for 24 h, respectively. LPA-E14 and HBPA-E14-1 were both with a weight ratio of 50:1 to mRNA. Samples were added at the final mRNA concentration of 0.125, 0.25, 0.5, 1, 2, 4 and 8 μg/mL per well. After 24 h, 10 μL of CCK8 detection reagent was added to each well, and absorbance at 450 nm was measured using a microplate reader.

### 7. *In vivo* mRNA transfection efficiency study

Finally, we tested the effect of HBPA-E14 on mRNA transfection *in vivo*. C57BL/6 mice (6 to 7 weeks old, about 20g) were raised at the SPF (specific pathogen-free) environment level. The in vivo-jetRNA/mRNA (in vivo-jetRNA), LPA-E14/LPA-PEG/mRNA (LPA/LPA-PEG), HBPA-E14-1/mRNA (HBPA), HBPA-E14-1/HBPA-PEG-1/mRNA (HBPA/HBPA-PEG) and Lyophilized HBPA-E14-1/HBPA-PEG-1/mRNA (Lyophilized HBPA/ HBPA-PEG) polyplexes were pulmonary administrated into the mice (using a pulmonary injection needle, 5 μg/mouse). After 24 h, the transfection efficiency of the luciferase mRNA in the body was measured as follows, using the saline as a blank control.

After mRNA transfection, the mice were euthanized, and lung tissues were harvested. Subsequently, the tissue was carefully placed into a grinding tube containing magnetic beads and subjected to grinding at 75 Hz with 4 cycles, each involving a 20 s running phase followed by a 20 s pause. after grinding, 20 μL of tissue grinding fluid was added into each well of the microplate reader, adding with 140 μL of assay buffer and thoroughly mixed. Then, 40 μL of D-Luciferin was added to the mixture and the luminescence was measured at 560 nm.

### 8. Statistical Analysis

Results were analyzed using GraphPad Prism (version 9.2.0 for Windows). Data are presented as mean ± SD. One- and two-way ANOVA with Dunnett’s multiple comparison test or Tukey’s multiple comparison were used where appropriate, unless otherwise indicated. Values were considered significantly different at p < 0.05.

